# Analysing the Yeast Complexome - The Complex Portal rising to the challenge

**DOI:** 10.1101/2020.11.03.367086

**Authors:** Birgit H M Meldal, Carles Pons, Livia Perfetto, Noemi Del-Toro, Edith Wong, Patrick Aloy, Henning Hermjakob, Sandra Orchard, Pablo Porras

## Abstract

The EMBL-EBI Complex Portal is a knowledgebase of macromolecular complexes providing persistent stable identifiers. Entries are linked to literature evidence and provide details of complex membership, function, structure and complex-specific Gene Ontology annotations. Data is freely available and downloadable in HUPO-PSI community standards and missing entries can be requested for curation. In collaboration with *Saccharomyces* Genome Database and UniProt, the yeast complexome, a compendium of all known heteromeric assemblies from the model organism *Saccharomyces cerevisiae*, was curated. This expansion of knowledge and scope has led to a 50% increase in curated complexes compared to the previously published dataset, CYC2008. The yeast complexome is used as a reference resource for the analysis of complexes from large-scale experiments. Our analysis showed that genes coding for proteins in complexes tend to have more genetic interactions, are co-expressed with more genes, are multifunctional, localize more often in the nucleus, and are more often involved in nucleic acid-related metabolic processes and processes where large machineries are the predominant functional drivers. A comparison to genetic interactions showed that about 40% of expanded co-complex pairs also have genetic interactions, suggesting strong functional links between complex members.

## Introduction

Many proteins exist as part of stable, macromolecular complexes that act as functional units in the cell. Identifying such complexes is crucial for a systems level understanding of biological processes. The EMBL-EBI Complex Portal (www.ebi.ac.uk/complexportal, (1, 2) is a manually curated, encyclopaedic resource of macromolecular complexes from a number of key model organisms, including *Saccharomyces cerevisiae*. Entries describe assemblies of two or more macromolecules (proteins, nucleic acids, small molecules) for which there is evidence (experimental or inferred) that these molecules stably interact with each other and have a demonstrated molecular function. Unlike other compendia of complexes, such as CORUM (3), it not only lists the protein composition of each complex but it also includes non-protein components, stoichiometry (when known), topology (including intra-complex binary interactions), and provides both, a free-text and structured description of complex function and properties (Figure 1). Each entry is linked to a range of related resources such as complex-centric Gene Ontology (GO) annotations (4, 5), structure determinations deposited in the wwPDB (6) or the role of the complex in a pathway in Reactome (human-only) (7). Links to these and other resources are provided both via cross-referencing and the integration of widgets on the website to display Reactome pathways diagrams, structures via the PDBe LiteMol App (8) and gene expression data via the Expression Atlas widget (9). Versioning of the stable accession numbers indicates when a complex has been significantly updated, for example by the addition or removal of a protein subunit from the list of participants. The data are freely available and downloadable in the HUPO-PSI community standard PSI-MI XML3.0 (10), MI-JSON and tab-delimited ComplexTab formats (2).

**Figure 1:**
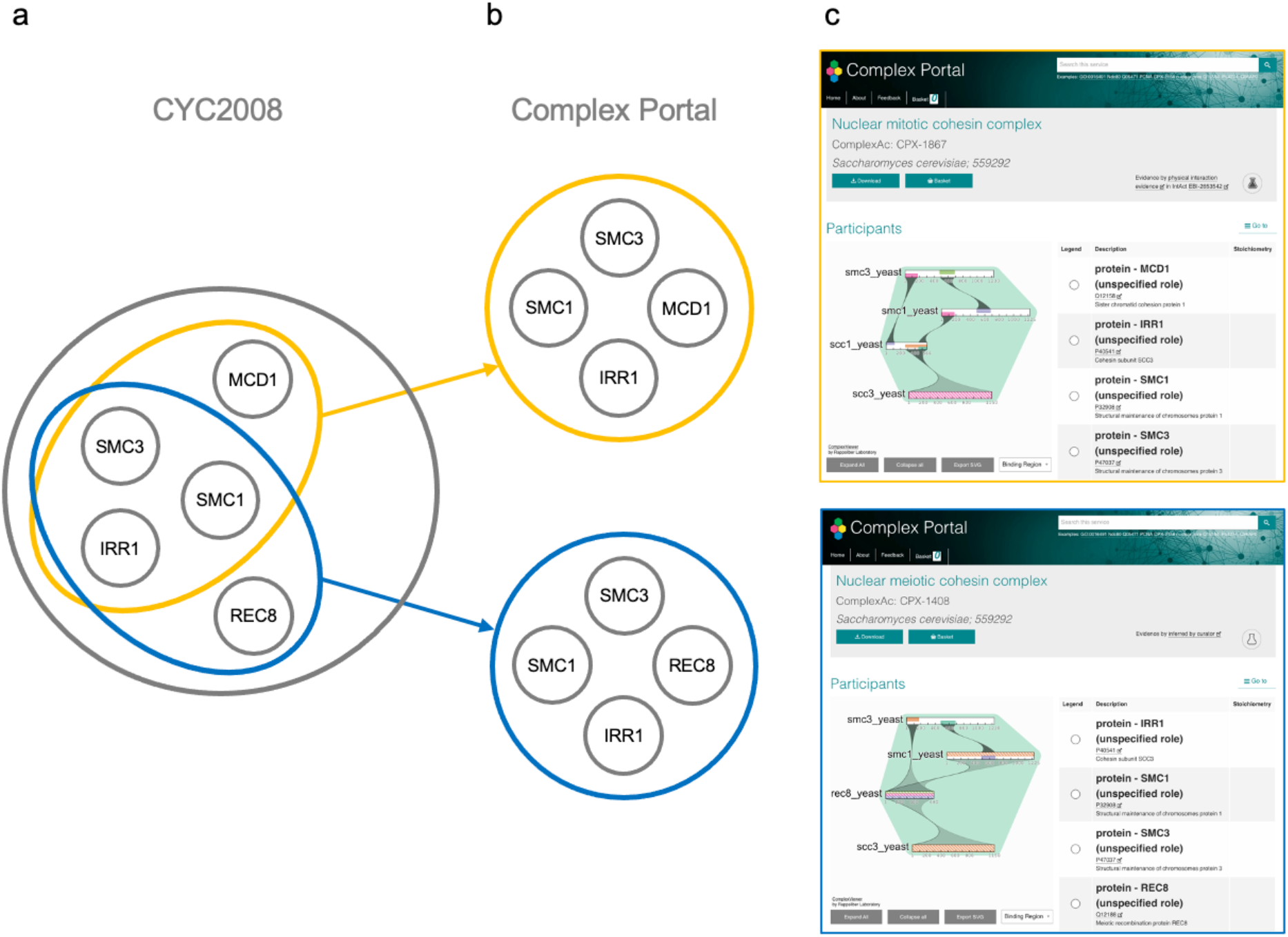
The nuclear cohesion complex is curated as one entry in CYC2008 (a) but represented by two, process-specific complexes in Complex Portal (b), one involved in mitosis and one involved in meiosis. The two complexes differ by one subunit. (c) Screenshots of the Details page of these complexes in Complex Portal.

*Saccharomyces cerevisiae* (henceforth referred to as “yeast”) is an important model organism for our understanding of the biology of all eukaryotic organisms and significant effort has gone into identifying all its stable complexes. However, until recently, information about such complexes was scattered across many publications and in different databases. An early effort to concatenate these data was the, now deprecated, MIPS yeast complex database (11). Domain-specific resources such as structural data in wwPDB, functional statements and Gene Ontology annotations on the protein pages of UniProt (12) and gene pages of the *Saccharomyces* Genome Database (SGD; www.yeastgenome.org) (13) provide highly-detailed, component-specific information only. It was very difficult to derive a picture of the complete yeast complexome without systematically integrating information from these and other sources. Additionally, molecular interaction databases such as those maintained by members of the IMEx Consortium (14) provide experimentally-derived interaction data without combining evidence from multiple sources for a whole complex. Several studies in the early 2000s predicted yeast complexes based on high-throughput yeast two-hybrid (15, 16) or affinity-purification methods (17–19) but only few studies included systematic validation by way of small-scale experiments and manual curation (20). In 2009, Pu *et al.* published a comprehensive analysis of 400 highly inter-connected assemblies derived from high-throughput experiments (<u>Y</u>east <u>H</u>igh <u>T</u>hrough<u>P</u>ut, YHTP2008) and also a compendium of 408 literature-derived, manually curated complexes based on small-scale experiments (<u>C</u>urated <u>Y</u>east <u>C</u>omplexes, CYC2008) (21). Whilst both datasets contained approximately 400 entries, less than 20% of these were identical to each other. However, in the 12 years since this set was first published, significant advances have been made in the field of interaction biology and considerably more high-quality datasets are now available to contribute to our understanding of this field. This has allowed a re-evaluation of the data and in 2018 the first version of an updated and enhanced dataset of known yeast complexes, the “yeast complexome”, was released in the Complex Portal. Additional complexes are being added to the dataset on an ongoing basis, if and when they are experimentally verified. In this paper we explore the yeast complexome and compare the extent and depth of data available through the Complex Portal to other resources that contain data on yeast complexes, namely to the curated and predicted complexes from Pu *et al.* and complexes predicted based on all experimental protein-protein interactions in the IntAct molecular interaction database (22). Compared to CYC2008, the Complex Portal dataset contains almost 50% more entries (589 vs 408), covers 4% more of the yeast proteome (30% vs 26%) and includes additional detail about the complexes as described above. Finally, we compare and contrast protein complex co-membership with the global genetic interaction network (23) and found that both datasets significantly overlap.

## Methods

### Source data for the Yeast Complexome

The data for the yeast complexome was derived from detailed literature searches and collated in collaboration with curators based at UniProt and SGD. A draft list of putative complexes was created based on the following sources: the CYC2008 dataset, UniProtKB SUBUNIT comment lines search with keywords “found in a complex with”, a close collaboration with SGD who provided a list of identified complexes and by directed literature searches. A complex is only included in the Complex Portal dataset if there is literature evidence for its existence and functional role *in vivo*. Complexes that were identified based only on either high or low-throughput analyses without the presence of further verification experiments or functional assays were not included. 13 homomers have been curated, to date, because the protein was also present in a related heteromeric complex. It should be noted that homomers have largely been omitted from manually curated datasets because it is often challenging to demonstrate experimentally if their function requires oligomerization and their generic functions are already described in the UniProtKB database. Literature searches and the collaboration with SGD are ongoing and new complexes are being added to the dataset when they are experimentally identified.

### The datasets

The protein complex datasets analysed were the following:

- Complex Portal - 589 complexes (release 228, 16/11/2019)
- CYC2008 - 408 manually-curated complexes (21)
- YHTP2008 - 400 predicted complexes (21)
- IntAct-LT - 332 predicted complexes derived from low-throughput experiments in IntAct (release 228, 16/11/2019)
- IntAct-HT - 689 predicted complexes derived from high-throughput experiments in IntAct (release 228, Nov 2019)

To enable direct comparison of protein complex components represented in the Complex Portal and IntAct, gene locus IDs in CYC2008 and YHTP2008 were mapped to UniProt ACs using the UniProt Mapping service web application (UniProt Release November 2019). Ambiguous mappings, where a locus could be mapped to more than one UniProt entry with the same sequence, were expanded to include all potential mapping pairs.

Complex Portal data was exported in ComplexTab format. Where complexes are part of larger assemblies (sub-complexes) these were expanded to provide a list of unique UniProtKB identifiers. Sets of paralogous ribosomal proteins were expanded to a full list, therefore all potential UniProtKB identifiers were included in the analyses. The expansion of paralogous proteins leads to an over-inflation of the subunit count per complex for the two ribosomal subunits but is the only way to include all proteins in the comparative analysis. As stoichiometry information is only available in a limited number of Complex Portal and IntAct entries and often missing due to a lack of available evidence, it was ignored and comparisons were based on unique protein identifiers only. Non-protein complex members such as nucleic acids and small molecules were not included as these are not provided in full by any resource other than the Complex Portal.

IntAct complexes were derived from all yeast-yeast interactions in IntAct release 228. Interactions were exported in MI-TAB2.7 format and split into those derived from papers with 100 or less interactions/paper and those with more than 100 interactions/paper. Complexes were predicted using the Cytoscape App ClusterONE (24) using default parameter settings, MI-score values as weights and a minimum cluster size of n=3.

### Functional analyses

For the selection of genetic interactions we used the global yeast genetic interaction network, the first comprehensive genetic interaction map in any organism (23). The network was constructed by evaluating the growth defects associated with the majority of the ~18 million possible gene pairs in yeast, and includes ~350,000 positive and ~550,000 negative genetic interactions. Non-essential genes were queried by deletion alleles and essential genes by temperature-sensitive and DAmP alleles. However, we disregarded the DAmP data because few DAmP alleles had an effect on cellular fitness. For pairs of genes screened more than once (for instance, pairs involving genes queried using different alleles) a consensus approach was implemented in which we considered a given pair to have a genetic interaction if that was the result in at least half of the screens.

Interacting protein pairs in a complex (i.e. co-complex pairs) were inferred by matrix expansion of all complexes. UniProt identifiers were mapped to ORFs in order to compare inferred physical interactions and genetic interactions as the latter are provided as ORFs. Background pairs (i.e. “no co-complex pairs”) were defined as those pairs of proteins present only in different complexes. The fractions of co-complex and background pairs with positive and negative interactions were calculated, considering only pairs of proteins whose genes were present in the genetic interaction network (52%, 51%, 55%, 58%, and 64% of co-complex pairs in CP, CYC, YHTP, IntAct-LT, and IntAct-HT, respectively). Statistical significance was calculated by Fisher’s exact tests.

In addition to genetic interactions, we evaluated the overlap of co-complex relationships with the co-expression, co-localization, and co-annotation functional standards. In all cases, only protein pairs for which functional data was available were considered. The co-expression standard was derived from the MEFIT co-expression network, which integrates data from multiple microarray datasets (25). Pairs with a MEFIT score >1.0 were considered to be co-expressed. The co-localization standard was based on a previous high-throughput study (26). Protein pairs localized in one or more shared cellular compartments were considered to be co-localized. The co-annotation standard is based on GO biological process annotations and disregards very frequently annotated GO terms as described in a previous work (23).

To obtain a comprehensive view of the differences between those proteins participating in complexes and those that do not, the following characteristics were compared: genetic interaction degree calculated on array genes and averaging estimates across the different alleles of a gene (23), co-expression degree calculated as the number of co-expression relationships per gene (see above), gene conservation in other species (27), expression variation (28), fitness of non-essential gene deletion alleles (23), PPI degree (from IntAct yeast-yeast interaction, release 234 (09/07/2020), restricted to high-throughput dataset with more than 100 interactions per publication as it reduces the bias from confirmatory small-scale experiments), multifunctionality of proteins based on the number of biological process annotations in GOSlim (downloaded from SGD, July 2020), fraction of disordered residues downloaded from d2p2.pro (29), being essential (30), being a gene duplicate defined as having a paralog in YeastMine (31), being a membrane protein (32) as well as subcellular localization (26) and broad functional classes (33). For each numerical feature, values were z-score normalized using the median and the standard deviation of the values for the background proteins. Statistical significance was evaluated using two-sided Mann-Whitney U tests. For each binary feature, fold enrichment was calculated as the ratio of complex members with that feature divided by the ratio of non-complex members with that feature. Statistical significance was calculated by two-sided Fisher’s exact tests.

The relative difference in transcript counts, expression variance, protein abundance, and protein halflife was calculated for co-complex and background pairs. For every pair and measure, we calculated the maximum (MAX) and minimum (MIN) value within the pair. The relative difference was then calculated as (MAX-MIN)/MAX. The larger this score is, the larger the difference between the pair of proteins/genes. Statistical significance was calculated using two-sided Mann-Whitney U tests.

Direct and indirect contacts were selected from a set of Complex Portal complexes of size 3 or larger that contained information for both types of contacts. Self interactions were ignored. Protein pairs belonging to different complexes of the selected set were defined as background. Genetic interaction profile similarity values were downloaded from http://thecellmap.org (34), considering both essential and non-essential genes, and averaging similarity values across alleles of the same gene.

A list of 12 high level GO terms (Table 2) was manually selected to best represent processes and functions related to nucleic acids as well as the component term “nucleus”. These terms were used to build a bespoke SLIM and all annotations to yeast proteins using these terms and their children were exported on 09/10/2020. This list of GO terms was used to filter all Complex Portal complexes that were also annotated to any of these terms. This analysis was only performed on the Complex Portal dataset as there are no complex-specific GO annotations for the other datasets.

**Table 2:** Number of Complex Portal complexes annotated to nuclear and nucleic acid related GO terms

| GO term ID | GO term name | GO class | Complexes |
| --- | --- | --- | --- |
| GO:0006139 | nucleobase-containing compound metabolic process | biological_process | 159 |
| GO:0010467 | gene expression | biological_process | 101 |
| GO:0051276 | chromosome organization | biological_process | 73 |
| GO:0006974 | cellular response to DNA damage stimulus | biological_process | 38 |
| GO:0071826 | ribonucleoprotein complex subunit organization | biological_process | 14 |
| GO:0006997 | nucleus organization | biological_process | 2 |
| GO:0005634 | nucleus | cellular_component | 197 |
| GO:0003676 | nucleic acid binding | molecular_function | 121 |
| GO:0140098 | catalytic activity, acting on RNA | molecular_function | 37 |
| GO:0140110 | transcription regulator activity | molecular_function | 16 |
| GO:0140097 | catalytic activity, acting on DNA | molecular_function | 15 |
| GO:0045182 | translation regulator activity | molecular_function | 12 |

### Analysis Tools

Data manipulation and visualisations were performed in R (data.table, splitstackshape, reticulate, rio, ggplot2, scales), Python and Excel. Unique versus shared sets of complexes were identified using Venny (https://bioinfogp.cnb.csic.es/tools/venny/).

## Results and Discussion

### The Yeast Complexome in the Complex Portal

*Saccharomyces cerevisiae* complexes were captured in the Complex Portal leading to yeast being the first completed species complexome. It is the largest manually-curated compendium of yeast multi-molecular complexes, comprising 589 complexes, 1930 proteins and 15,863 co-complex relationships. In order to identify all known yeast complexes we gathered information from a number of sources (CYC2008 complexes, UniProt, SGD, literature publications). Some complexes that are included in these sources have not been included in the Complex Portal because they have since been identified as part of a bigger complex or they lack clear experimental evidence for their existence and their functional role *in vivo*. These putative complexes are kept in a separate list and are periodically revisited to see if more evidence has been published. Collaborations with SGD are ongoing and we update existing entries and add new ones when new evidence comes to light.

Compared to other resources, the Complex Portal provides added value through its greater scope of annotation. Each complex entry has a manually annotated description of their function and physical properties and includes stoichiometry and topological information when available. The Evidence and Conclusion Ontology (ECO) (35) is used to indicate the type of evidence we have for each entry and where interaction evidence is available in an IMEx member database, the wwPDB or EMDB (36) cross-references are provided. Each complex is annotated to GO terms specific for the complex and a selection of supporting literature references are provided. Versioning allows easy tracking of changes in complex composition. Additionally, the data is downloadable in three different community standard formats and as a live resource it gets updated every two months.

### Dataset comparisons

The yeast complex dataset published in the Complex Portal is the first manually annotated yeast complex dataset since the publication of CYC2008 by Pu *et al.* in 2009. We compare these two manually curated datasets with each other and with corresponding experimentally-derived predicted complexes from YHTP2008 and IntAct release 228 (16/11/2019). The IntAct data was split into low and high throughput publications setting a cut-off at 100 interactions per publication. See Table 1 for a summary of the five datasets and Figure 2 for the distribution of unique proteins per complex.

**Table 1:**
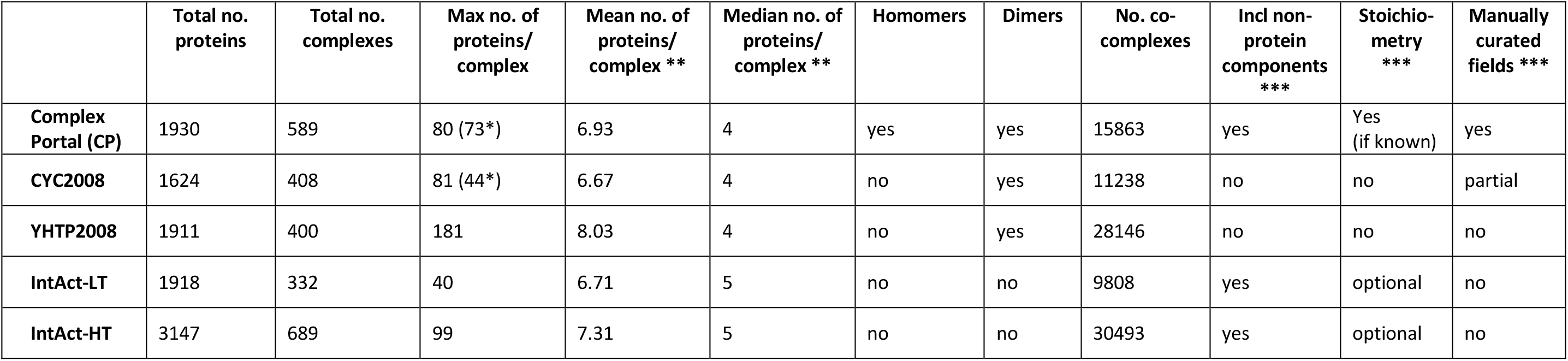
Basic statistics about the five complex datasets

|  | Total no. proteins | Total no. complexes | Max no. of proteins/complex | Mean no. of proteins/complex ** | Median no. of proteins/complex ** | Homomers | Dimers | No. co-complexes | Incl non-protein components *** | Stoichiometry *** | Manually curated fields *** |
| --- | --- | --- | --- | --- | --- | --- | --- | --- | --- | --- | --- |
| **Complex Portal (CP)** | 1930 | 589 | 80 (73*) | 6.93 | 4 | yes | yes | 15863 | yes | Yes (if known) | yes |
| **CYC2008** | 1624 | 408 | 81 (44*) | 6.67 | 4 | no | yes | 11238 | no | no | partial |
| **YHTP2008** | 1911 | 400 | 181 | 8.03 | 4 | no | yes | 28146 | no | no | no |
| **IntAct-LT** | 1918 | 332 | 40 | 6.71 | 5 | no | no | 9808 | yes | optional | no |
| **IntAct-HT** | 3147 | 689 | 99 | 7.31 | 5 | no | no | 30493 | yes | optional | no |

**Figure 2:**
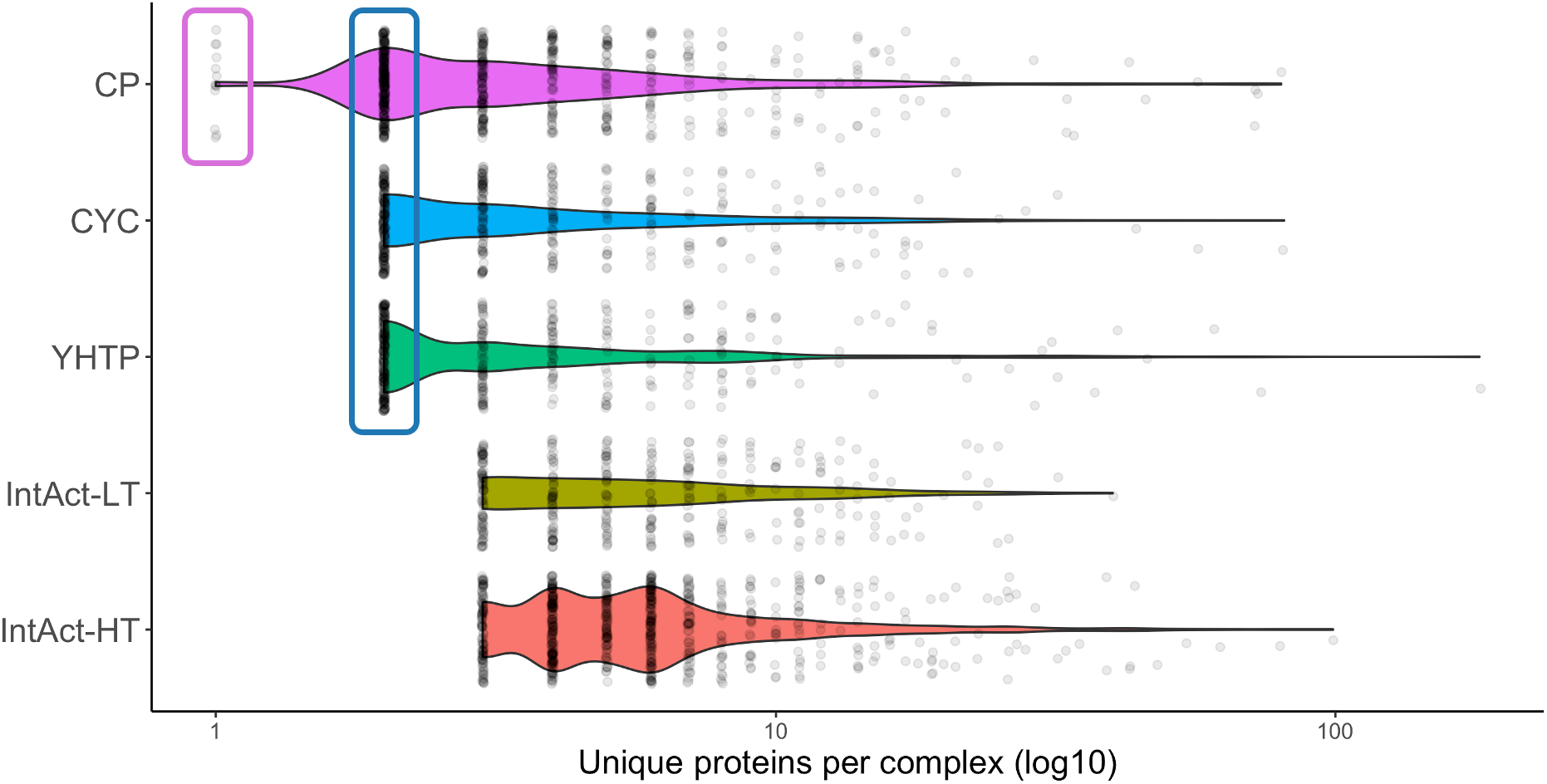
Distribution of number of unique proteins per complex. Homomers are found in the pink rectangle and heterodimers in the blue rectangle. Total number of complexes per dataset: CP = 589, CYC = 408, YHTP = 400, IntAct-LT = 332, IntAct-HT = 689

The two manually-curated datasets share 1543 proteins (80% and 95%, respectively): 387 proteins are unique to the Complex Portal and 81 unique to CYC2008 (Table 1, Figure 3a); overall, Complex Portal and CYC2008 complexes cover 32% and 27% of the yeast proteome, respectively. The reason for the relatively low proteome coverage may be multifaceted: both datasets have concentrated on stable, macromolecular machines whereas many proteins may be found in more transient interactions, such as signalling assemblies or enzyme-substrate interactions. The identification of protein complexes may also be limited by technological constraints and some complexes simply cannot be purified by existing methods, for example insoluble membrane components.

**Figure 3:**
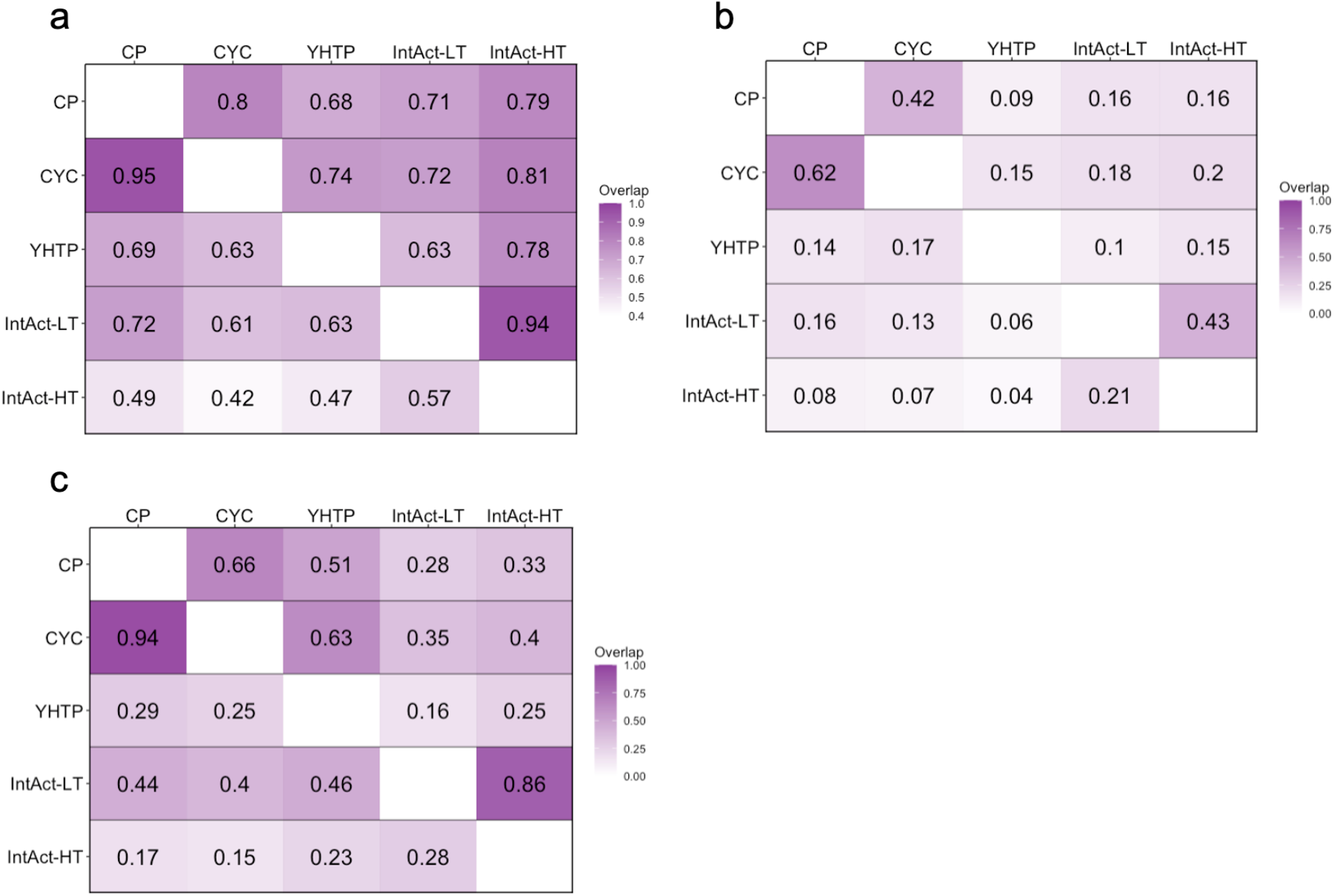
Fraction of (a) proteins (CP = 1930, CYC = 1624, YHTP = 1911, IntAct-LT = 1918, IntAct-HT = 3147), (b) complexes, based on Jaccard Index = 1.0 for complexes with a minimum of 3 protein participants (CP = 345, CYC = 236, YHTP = 208, IntAct-LT = 332, IntAct-HT = 689) and (c) co-complex pairs shared between two any datasets (CP = 15863, CYC = 11238, YHTP = 28146, IntAct-LT = 9808, IntAct-HT = 30493). Each row compares the overlap of both datasets to the total number of entities in the dataset given on the left.

The Complex Portal contains 589 yeast complexes compared to 408 in the CYC2008 dataset, a 44% increase (Table 1). They share 286 identical complexes which responds to 49% of Complex Portal complexes and 70% of CYC2008 complexes (Jaccard Index = 1.0) (Figure 4a). When reducing protein identity matching to a minimum of 50% (Jaccard Index = 0.5) the overlap is over 80% for both datasets (Figure 4c). There are many more complexes in the Complex Portal than in CYC2008 because a large amount of knowledge has accumulated in the intervening 12 years. On the other hand, approximately 30 CYC2008 complexes were not re-curated into the Complex Portal because the available interaction evidence does not meet current curation criteria (1) or because they are now believed to be part of larger complexes. These complexes remain on a watch list and will be added if sufficient evidence becomes available. Complex Portal complexes also contain 94% of CYC2008 co-complex pairs while CYC2008 complexes only contain 66% of Complex Portal co-complex pairs (Figure 3c).

**Figure 4:**
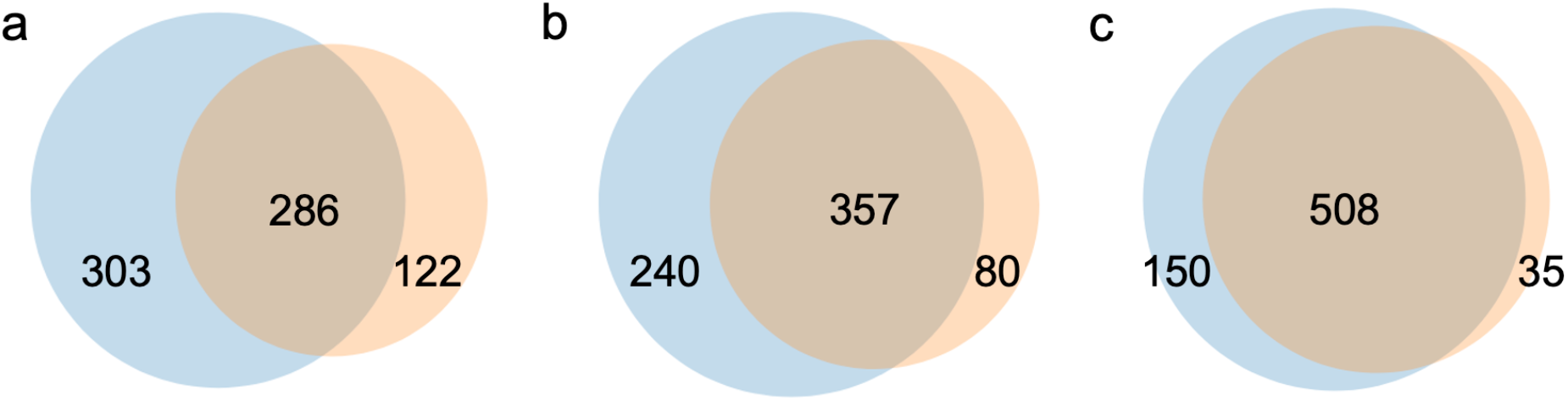
Overlap of complexes by protein identities and decreasing stringencies for complex membership between Complex Portal (n = 589, blue) and CYC (n = 408, orange). (a) JI = 1.0, (b) JI = 0.75, (c) JI = 0.5. JI = Jaccard Index. NB: Total numbers per dataset for JI = 0.75 and JI = 0.5 are higher than the absolute number of complexes per dataset as one complex can be broken down into more than one partial complex that matches a complex in the other dataset.

The IntAct yeast interactome contains a total of 124,918 yeast-yeast binary interactions containing 5850 unique proteins or 97% of the yeast proteome (proteome = 6049 proteins) and 18 interactions between a yeast protein and a yeast complex. A topological clustering analysis of the IntAct yeast interactome was performed using the Cytoscape App ClusterONE, restricting accepted clusters to those with 3 or more proteins. The resulting clusters encompassed only just over half the proteome (3280 proteins, 54%) and predicted 332 complexes from low-throughput publications (IntAct-LT) and 689 complexes from high throughput publications (IntAct-HT) (Table 1). Only a third of the proteome was present in the 400 YHTP2008 predicted complexes based on high throughput data (1911 proteins, 32%).

Complex sizes (Figure 2) are difficult to compare as the minimum sizes are determined by the curation strategies (see Table 1 for a reference of which datasets contain homomers and dimers) and the maximum sizes determined by the handling of paralogous proteins. Where possible, Complex Portal curates separate complexes for each paralogous protein but in the case of the ribosomal subunits it creates sets for each paralogous pair. Similarly, CYC2008 often includes each paralogous gene locus in the same complex. The inclusion of paralogous proteins or loci in a complex artificially inflates its maximum (and with that the mean and median) size. Likewise, clustering algorithms tend to group paralogous proteins together. Therefore, the largest complexes are found in the predicted datasets of YHTP2008 and IntAct-HT. Excluding the ribosomal subunits that contain multiple paralogous pairs of proteins or loci, the maximum size of a complex in the Complex Portal is 73 and in CYC2008 is 44.

However, despite the issues with minimum and maximum complex sizes, the overall complex size distributions are very similar. The majority of complexes contain 10 or fewer unique proteins with a rapidly reducing tail. This is dataset-independent and demonstrates that most proteins function within a relatively small group of partners. There are a few larger complexes in the Complex Portal than in CYC2008. ClusterOne predicts no complexes larger than 40 proteins/complex for the IntAct-LT dataset resulting in the smallest complex size distribution of all datasets. In comparison, IntAct-HT has the highest predicted complex size distribution of all datasets when ignoring the expanded ribosomal complexes. The IntAct-HT dataset includes many affinity purification experiments, which can identify large associations of co-purifying proteins which in turn result in more centralized and heavily-connected areas of the underlying interactome. Such heavily-connected areas in the interactome result in many overlapping clusters that have a tendency to get combined into superclusters by the ClusterOne algorithm.

We also compared the manually-curated complexes with those predicted from experimental protein-protein interaction (PPI) evidence. The overlap between any curated and predicted dataset never exceeded 20% in any comparison with a Jaccard Index of 1.0 (Figure 3b). The IntAct-HT complexes contain an even smaller overlap with either of the curated complex datasets (7-8%) than the IntAct-LT or YHTP2008 complexes (13-17%). At the protein level, only 42-72% of proteins from an experimental dataset could also be found in a curated complex dataset while 68-81% of proteins in the curated datasets are also found in the experimental datasets (Figure 3a).

The low level of overlap between manually-curated and predicted complexes may be the result of a combination of factors: Firstly, experimentally-derived interactomes contain a lot more proteins than the complex datasets but incorporate fewer validated evidence than the often thoroughly and even functionally validated interaction evidence used to define curated complexes. Secondly, the need for a reductionist representation of the interactome, where multiprotein associations are reduced to binary pairs via spoke expansion methods introduces a bias in the internal topology of PPI evidence networks, potentially generating spurious associations. Finally, prediction algorithms are restricted to predicting heteromers and ClusterOne restricts clusters to size 3 and larger; therefore, any heterodimeric complexes are not included in the predicted datasets and were removed from the above comparisons for the overlap of complexes between the five datasets.

### Features of protein complexes, their proteins and genes that code for them

The properties of protein complex members were characterised using a panel of numerical and binary features (Supplementary Figure 1). Genes coding for proteins in complexes tended to have more genetic interactions and to be co-expressed with more genes. They were also more likely to be multifunctional, conserved across species, and present more stable expression patterns. Additionally, they often coded for proteins with a higher percentage of disorder, higher PPI degree and were enriched for essential genes and non-essential genes with larger fitness defects. On the other hand, these genes were depleted for duplicates and were less likely to code for membrane proteins. Localization patterns changed slightly across datasets. Proteins in complexes tended to localize more often in the nucleus and the nucleolus than other proteins, while they were less likely to be found in the vacuole. To further explore this finding, complexes in the Complex Portal dataset were analyzed for annotations to nuclear and nucleic acid-related processes and functions (Table 2) taking advantage of the complex-specific GO annotations available for this dataset. More than half of complexes (304/589, 52%) are annotated to at least one of these 12 selected terms or their children (Supplementary Table 2). 65% of these complexes (197/304) are annotated to “GO:0005634 nucleus” or a child term and 52% (159/304) to “GO:0006139 nucleobase-containing compound metabolic process” or a child term. In all datasets, proteins found in complexes were also significantly over-represented in processes where large machineries are the predominant functional drivers such as replication, transcription, translation, and ER to Golgi and trans-Golgi transports. This reflects how such processes require a variety of tightly regulated multimolecular machineries whose diversity has been thoroughly explored in the literature. However, proteins found in complexes were underrepresented in many signaling, transportation, and localization processes that are more often driven by single proteins. Importantly, most results were consistent across all five complex datasets.

### Multifunctionality

More than 70% of proteins in each dataset are only found in a single complex (Supplementary Figure 2) and there is no difference in this distribution between curated and predicted complexes. Only a few proteins from each dataset are found in two to five different complexes while CYC contains only a few and YHTP and IntAct-LT contain no proteins that occur in more than six complexes. Most of the proteins found in more than one complex are core subunits of complexes of which many different variants exist, such as cyclin-dependent kinases or ubiquitin ligases. In a recent analysis of datasets from several yeast interactome datasets it was demonstrated that this long right-hand tail of a few proteins occurring in many complexes is almost always significantly different from a random distribution (37). The random distribution estimates that proteins should be found in a maximum of 6-9 complexes while in the real data some proteins occur in >20 complexes, matching our observations.

There are 5 proteins that are found in *≥*4 complexes in the Complex Portal where the complexes are annotated to two or more unrelated pathways or complexes and three of these proteins are also found in more than one subcellular location when part of multiple complexes. Four of these proteins, H4 (P02309), LTV1 (P34078), SKP1 (P52286) and TAF14 (P35189), are regulatory subunits and one, PP12 (P32598), is a protein phosphatase (Supplementary Table 1). These five proteins have a relatively higher number of GO SLIM annotations compared to the rest (p < 0.0005, Supplementary Figure 3). All other complexes that share proteins are functionally-related homologues.

### Biological assessment of complexes via omics data

Genetic interactions identify combinations of genes that yield unexpected phenotypes when simultaneously mutated. Negative genetic interactions identify cases with more severe phenotypes than expected given the individual mutant phenotypes, whereas in positive genetic interactions the resulting phenotype is healthier. Both types of genetic interactions are a powerful tool for the characterization of genes and to elucidate the functional wiring of the cell (38).

Since genetic interactions identify potentially functional relationships between genes, we evaluated whether gene pairs coding for proteins within the same complex were enriched in genetic interactions using the global genetic interaction network (23). Genetic interactions have been explored for ~52% of the co-complex pairs defined in the Complex Portal dataset. Of these, 30% and 10% of genes coding for co-complex pairs had negative and positive genetic interactions, respectively. These represent a 4.4 and 2.4 fold increase, respectively, over what was observed in background pairs, i.e. pairs of genes coding for proteins in different complexes (p<0.05, Figure 5, Supplementary Figure 4). The significant overlap between genetic interactions and co-complex relationships is in agreement with previous studies (23, 39). This result was consistent across the different complex datasets, but the curated datasets and IntAct-LT showed a higher overlap with genetic interactions. A lower overlap of the high-throughput datasets, IntAct-HT and YHTP, with genetic interactions could be due to a larger fraction of indirect physical associations identified in weakly connected, large complexes in such studies. We found similar trends when comparing co-complex pairs to co-expression, co-localization, and co-annotation datasets (Figure 6, Supplementary Figure 5). In all cases, co-complex pairs had a higher overlap with these functional standards than background pairs and this overlap was particularly pertinent in the curated datasets. For instance, ~90% of co-complex pairs in the curated datasets were co-expressed, whereas the overlap for the remaining datasets ranged from 41% to 76%. Additionally, we observed more similar transcript counts, expression variance, and protein abundance and halflife for co-complex pairs than background pairs (Figure 7, Supplementary Figure 6), which reflects that members of the same protein complex tend to exhibit similar regulation patterns at a gene and protein level in order to act as a single coordinated biological unit.

**Figure 5:**
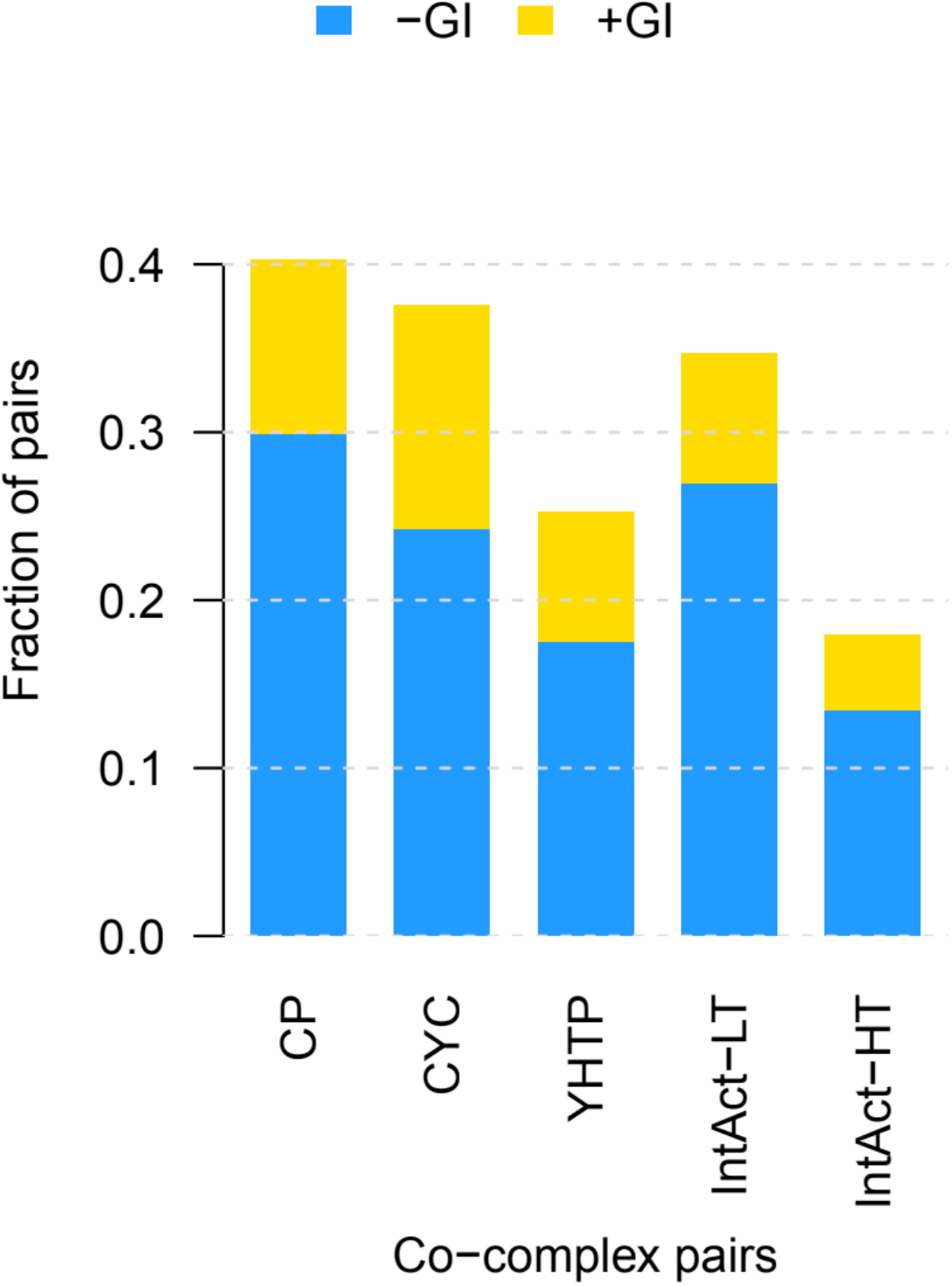
Fraction of co-complex pairs from each complex dataset that overlaps with negative (blue) and positive (yellow) genetic interactions.

**Figure 6:**
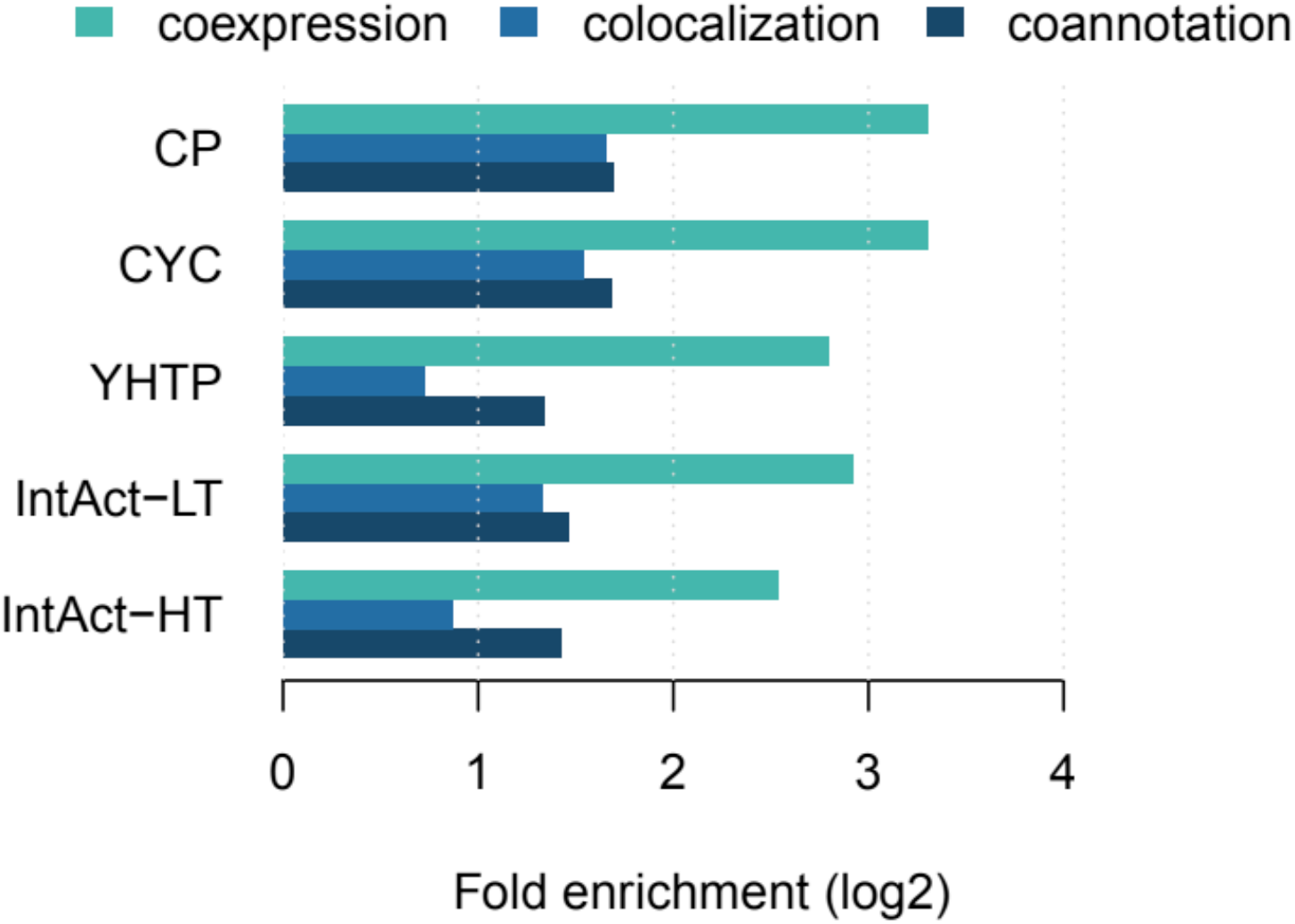
Fold enrichment of co-complex pairs compared to background pairs from all five datasets for co-expression, co-localization and co-annotation. All enrichments are statistically significant (p<0.05)

**Figure 7:**
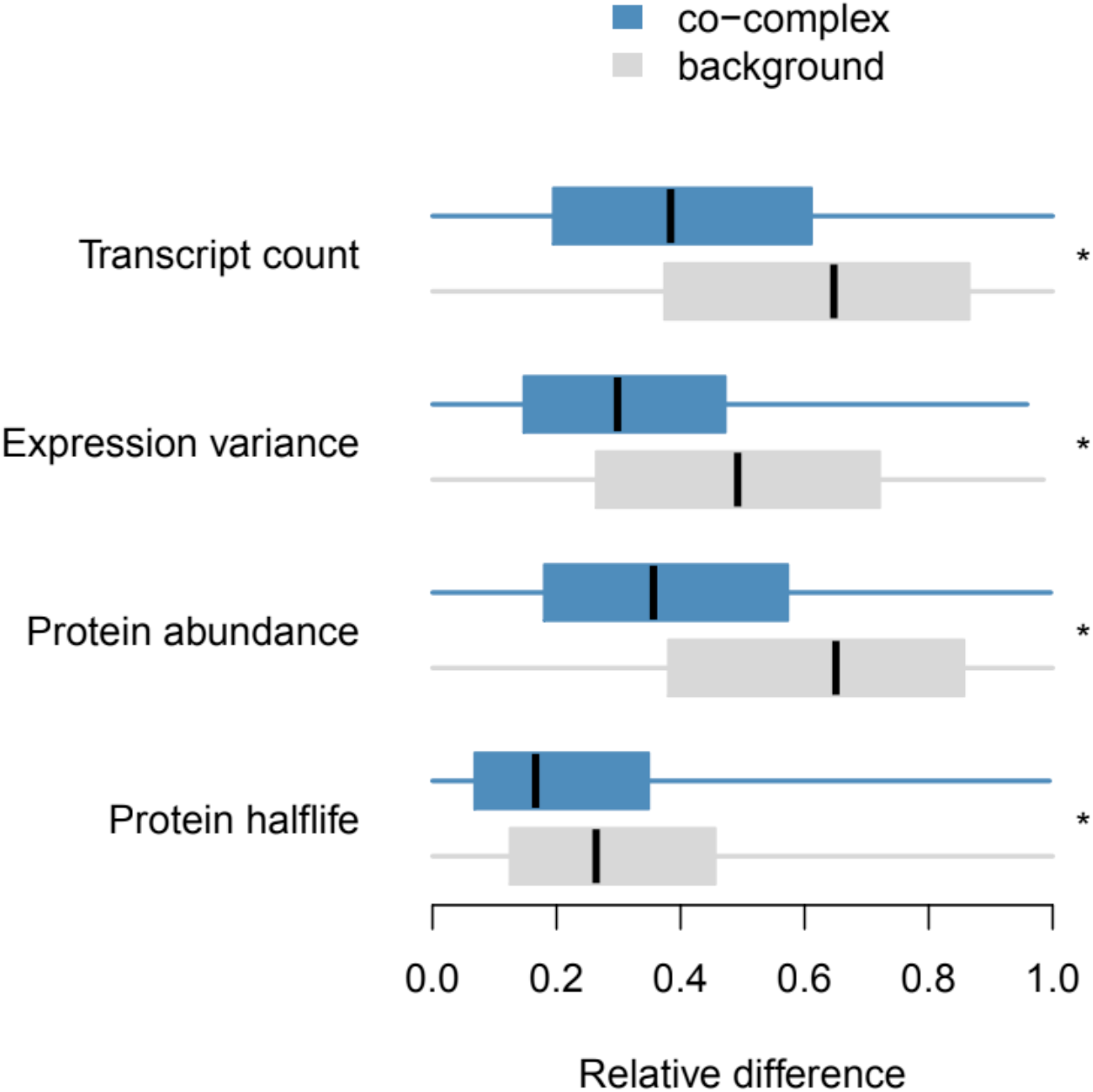
Relative difference in transcript counts, expression variance, protein abundance, and protein halflife for co-complex and background pairs in the Complex Portal. * p < 0.05.

Identifying the direct physical contacts within protein complexes can reveal sub-complex modules, improve the characterization of protein function, and help to interpret how mutations affect the phenotype. The Complex Portal is the only dataset that describes the internal connectivity of complexes, with detailed information for 237 complexes that have 3 or more participants. The functional relevance of this data was evaluated by comparing genetic interaction profiles (i.e., the set of genetic interactions of a gene) of direct and indirect contacts within protein complexes. These profiles are quantitative phenotypic signatures and revealed a higher similarity for gene pairs coding for proteins in direct contact (Figure 8; p<0.01 for all pairwise comparisons). This suggests that, in protein complexes with unknown internal connectivity, the analysis of genetic interaction profiles of the individual components may discriminate direct from indirect contacts.

**Figure 8:**
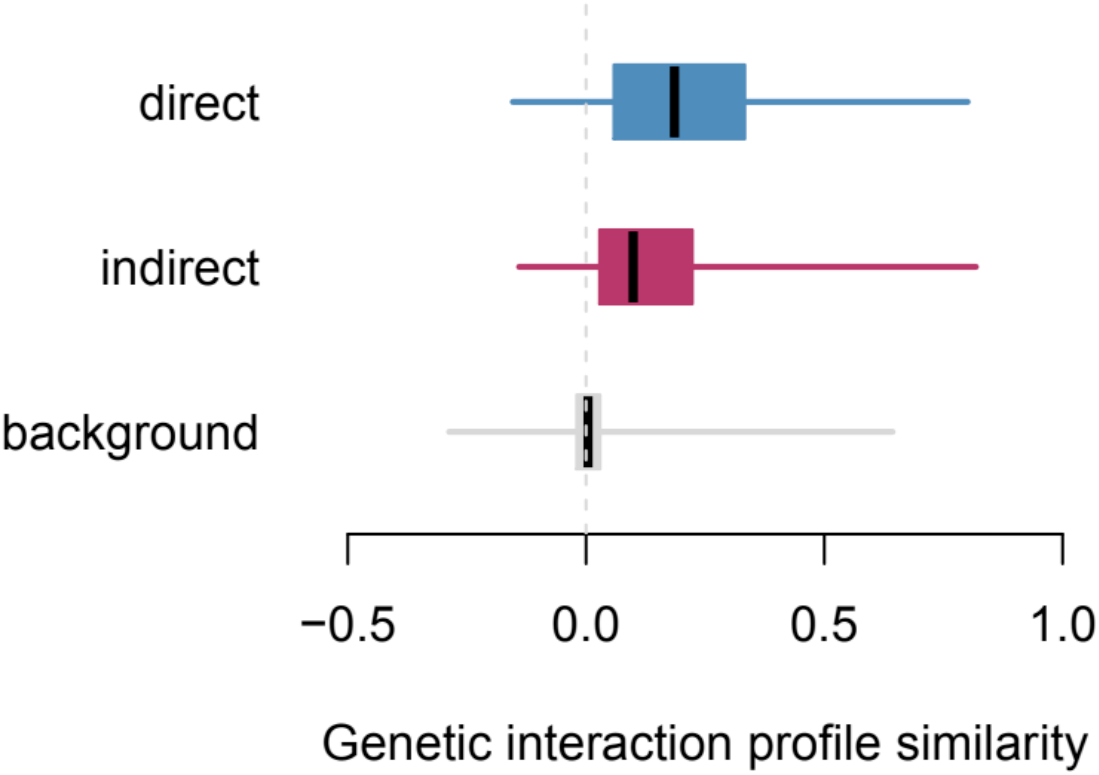
Genetic interaction profile similarities of gene pairs coding for proteins: in direct physical contact (blue), in the same complex but not in contact (red), in different complexes (grey). Boxes represent second and third quartiles, whisker first and forth. Horizontal lines in boxes represent the medians. All pairwise comparisons: p < 0.05

## Conclusions

Our knowledge of the biology of *Saccharomyces cerevisiae* has substantially improved over the last 12 years. The Complex Portal now provides almost 50% more complexes than did the previous compendium, CYC2008 (21), and these include more protein components, details on non-protein participants and more complex variants. The Complex Portal also provides a searchable website, a web service and three download formats.

Our set of curated yeast complexes shows a large overlap with previous curation efforts (i.e. CYC2008). However, these show a poor overlap when compared to predicted complexes. This may be due to large-scale affinity purification data producing clusters of apparently highly connected proteins as well as the presence of transient interactions in these datasets. This poor overlap also highlights that experimental protein-protein interactomes are a limited predictor for functional complexes which highlights the continuing need for a manually curated complex database.

Most proteins are found in only one complex and those found in two or more complexes tend to have the same function in multiple complexes. Only five proteins found in four or more complexes are linked to different processes showing that protein function is fairly conserved when they are part of complexes.

We highlight that there is a relative enrichment of multi-molecular machines in the nucleus and the nucleolus. These complexes are often involved in nucleic acid-related metabolic processes like replication, transcription and translation, plus other processes where multi-molecular assemblies are the predominant functional drivers such as ER to Golgi and trans-Golgi transports.

We found that the co-complex pairs overlap significantly with genetic interaction, co-expression, co-localization, and co-annotation datasets, which highlights the functional relevance of co-complex membership and the potential of protein complex datasets to address questions of biological interest. Members of the same co-complex also tended to present more similar regulation patterns which reflects the role of the protein complex as a coordinated biological unit. Genes coding for co-complex pairs in physical contact exhibited more similar patterns of genetic interactions, illustrating that the structural organization within complexes is key to interpret the results of functional studies. Importantly, contact information within complexes is only available in Complex Portal and not in the other complex datasets.

To date, the Complex Portal yeast complexome has been used to validate complexes in several large-scale studies (37, 39–42) and our stable identifiers are used as annotation objects and cross-references in several other curated databases, such as IMEx consortium partners, Gene Ontology (43), Genome Properties (44), MatrixDB (45), SGD (13), Reactome, Signor (46, 47) and Wikipathways (48) while other collaborations are under development, e.g. with PDBe (49). As we move to complete more complexomes, for example that of *Escherichia coli*, and continually improve our coverage of the human and mouse complexes, it will also be possible to improve our understanding of the evolution of these assemblies (50), and from there how the regulation of cellular processes has developed as organisms evolve.

We have shown how the Complex Portal yeast complexome is a key resource that significantly extends previously-available datasets. Our commitment to keep it updated and freely accessible ensures the scientific community can count on a stable, high-quality reference set for the study of multi-molecular machineries in yeast and other organisms.

We encourage our users to get in touch via the website if they find missing complexes or have suggestions on how to improve or extend our service.

## Data Availability

The complete yeast complexome is available for download from www.ebi.ac.uk/complexportal/download, the CYC2008 and YHTP2008 data from http://wodaklab.org/cyc2008/downloads and all files listing complexes and co-complexes used as input for our analyses have been deposited in Zenodo (10.5281/zenodo.4160609).

## Acknowledgements

We would like to thank Eliot Ragueneau for his help with figure design and Colin Combe for his contributions to the development and maintenance of ComplexViewer.

## Supplementary Data

Supplementary Table 1: Associated functions, process and complexes of proteins that occur in ≥4 complexes where the protein in question is not a core functional subunit of a set of paralogous complexes.

Supplementary Table 2: List of all Complex Portal complexes and their annotations to nuclear and nucleic acid related GO terms

Supplementary Figure 1: Gene and protein features of complex members compared to non-complex members (background). (a) Panel of numerical features. Yellow and blue dots identify features with significantly higher and lower values for complex members, respectively. Numerical values were z-score normalized using the median and the standard deviation of the background proteins. Dot size is proportional to the median z-score value of the proteins in complexes. (b-d) Panel of binary features (b), localization patterns (c), and functional classes (d). Fold enrichment for a particular binary feature was calculated as the ratio of complex members with that feature divided by the ratio of non-complex members with that feature. Yellow and blue dots identify features with significantly higher and lower ratios for complex members, respectively. Dot size is proportional to the fold enrichment.

Supplementary Figure 2: Multifunctionality of proteins in complexes in all five datasets (CP = 589, CYC = 408, YHTP = 400, IntAct-LT = 332, IntAct-HT = 689).

Supplementary Figure 3: Density distribution of GO SLIM biological process annotations in SGD for proteins found in Complex Portal complexes versus those not found in complexes. The number of annotations for 5 multifunctional proteins found in ≥4 complexes and annotated to two or more unrelated pathways or complexes are indicated with arrows.

Supplementary Figure 4: Fraction of protein pairs from each complex dataset that do not occur in the same complex (= background pairs) that overlaps with negative (blue) and positive (yellow) genetic interactions.

Supplementary Figure 5: Fraction of co-complex pairs (bars) and background pairs (white crosses on bars) of all five datasets that overlap with the co-expression, co-localization and GO co-annotation functional standards.

Supplementary Figure 6: Relative difference in transcript counts, expression variance, protein abundance, and protein halflife for co-complex and background pairs in CYC, YHTP, Intact-LT, and Intact-HT. * p < 0.05.

