## Supplementary material for "Analysing the Yeast Complexome - The Complex Portal rising to the challenge": Supplemetary Table 1 and Supplementary Figures 1-6

Supplementary Materials

Supplementary Table 1: Associated functions, process and complexes of proteins that occur in ≥4 complexes where the protein in question is not a core functional subunit of a set of paralogous complexes.

| **UniProt AC** | **UniProt name** | **Protein name** | **Gene name** | **Complex functions or processes** | **N complexes** |
|---|---|---|---|---|---|
| P52286 | SKP1_YEAST | Suppressor of kinetochore protein 1 | SKP1 | in SCF ubiquitin ligases, centromer complexes, regulation of vacuolar ATPase complex assembly | 13 |
| P32598 | PP12_YEAST | Serine/threonine-protein phosphatase PP1-2 | GLC7 | involved in regulation of endonucease activity, cell polarity/budding, spindle attachment and glucose homeostasis | 8 |
| P02309 | H4_YEAST | Histone H4 | HHF1 | pos & neg regulation of transcription, part of nucleosome | 7 |
| P35189 | TAF14_YEAST | Transcription initiation factor TFIID subunit 14 | TAF14 | part of several transcription factor complexes, chromatin remodelers and histone acetyltransferase | 5 |
| P34078 | LTV1_YEAST | Low-temperature viability protein 1 | LTV1 | processome variants and endosome cargo recycling | 4 |

Supplementary Table 2: List of all Complex Portal complexes and their annotations to nuclear and nucleic acid related GO terms

*As separate Excel file.*

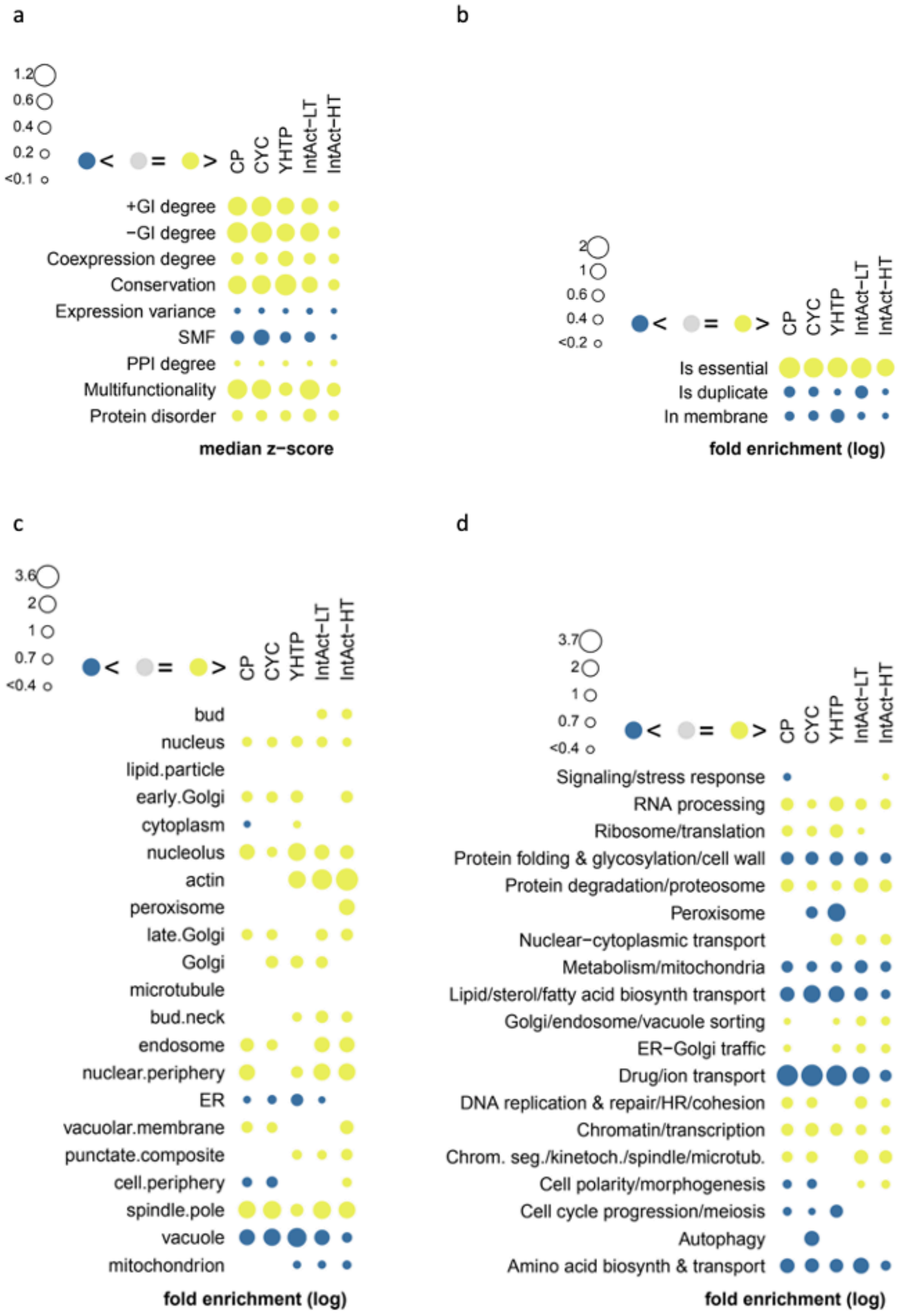


Supplementary Figure 1: Gene and protein features of complex members compared to non-complex members (background). (a) Panel of numerical features. Yellow and blue dots identify features with significantly higher and lower values for complex members, respectively. Numerical values were z-score normalized using the median and the standard deviation of the background proteins. Dot size is proportional to the median z-score value of the proteins in complexes. (b-d) Panel of binary features (b), localization patterns (c), and functional classes (d). Fold enrichment for a particular binary feature was calculated as the ratio of complex members with that feature divided by the ratio of non-complex members with that feature. Yellow and blue dots identify features with significantly higher and lower ratios for complex members, respectively. Dot size is proportional to the fold enrichment.

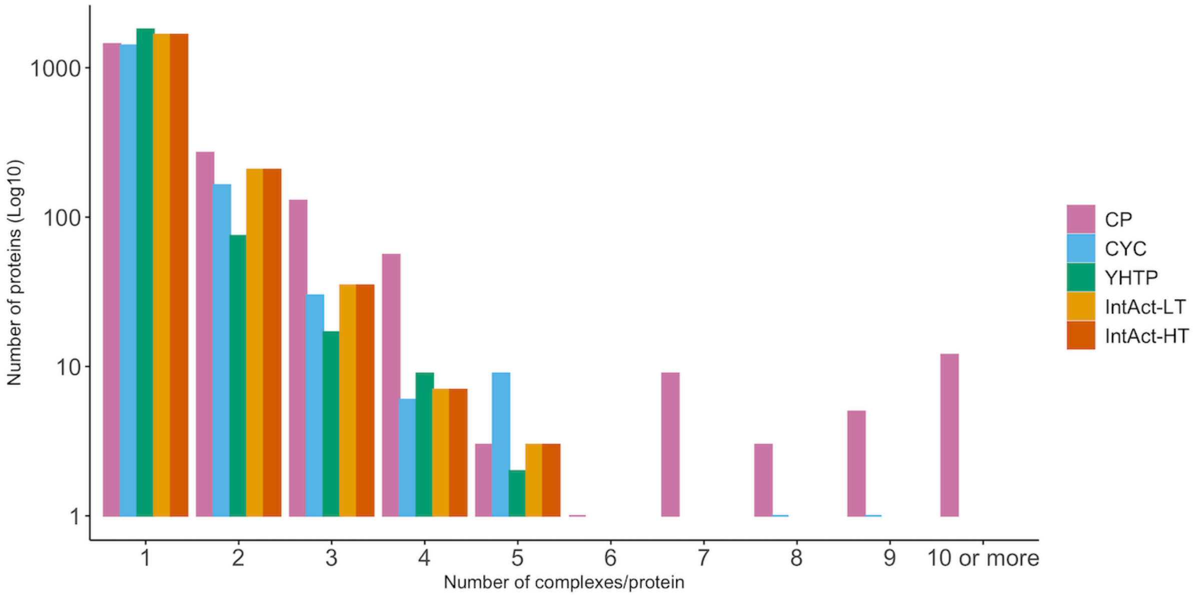


Supplementary Figure 2: Multifunctionality of proteins in complexes in all five datasets (CP = 589, CYC = 408, YHTP = 400, IntAct-LT = 332, IntAct-HT = 689).

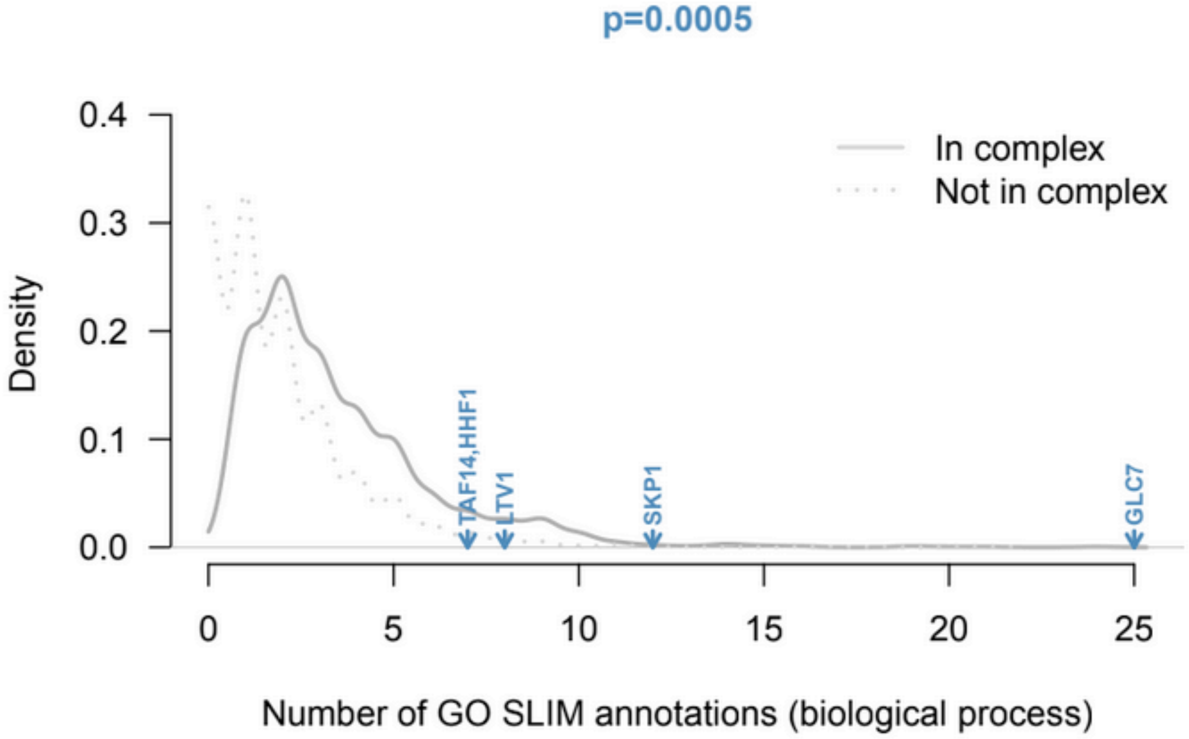


Supplementary Figure 3: Density distribution of GO SLIM biological process annotations in SGD for proteins found in Complex Portal complexes versus those not found in complexes. The number of annotations for 5 multifunctional proteins found in ≥4 complexes and annotated to two or more unrelated pathways or complexes are indicated with arrows.

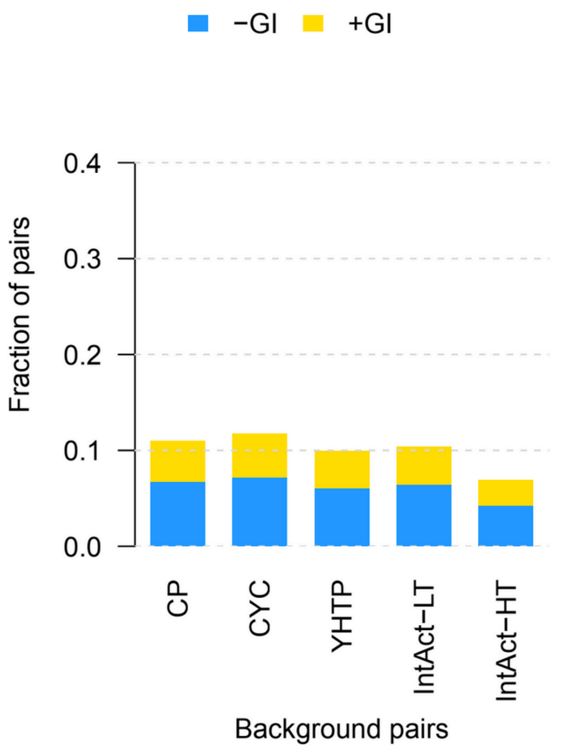


Supplementary Figure 4: Fraction of protein pairs from each complex dataset that do not occur in the same complex (= background pairs) that overlaps with negative (blue) and positive (yellow) genetic interactions.

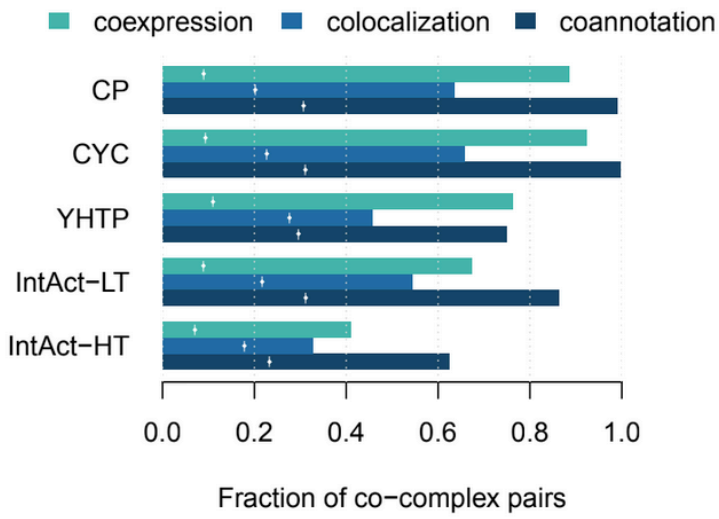


Supplementary Figure 5: Fraction of co-complex pairs (bars) and background pairs (white crosses on bars) of all five datasets that overlap with the co-expression, co-localization and GO co-annotation functional standards.

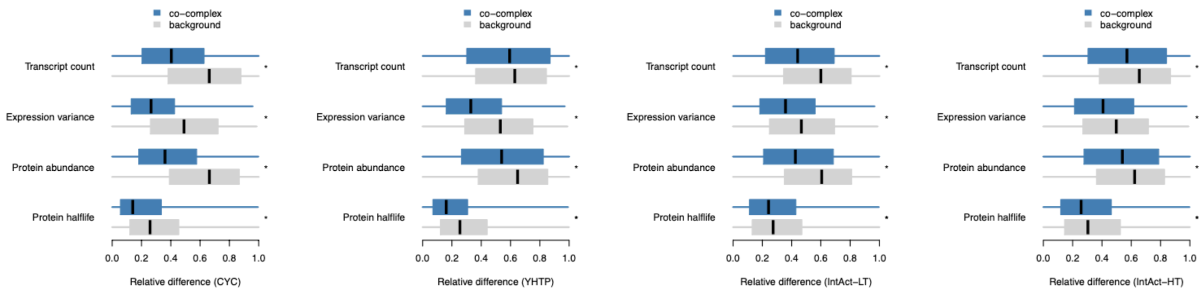


Supplementary Figure 6: Relative difference in transcript counts, expression variance, protein abundance, and protein halflife for co-complex and background pairs in CYC, YHTP, Intact-LT, and Intact-HT. * $p < 0.05$.
